# Inhibition of KDM1A activity restores adult neurogenesis and improves hippocampal memory in a mouse model of Kabuki syndrome

**DOI:** 10.1101/2020.03.11.986976

**Authors:** Li Zhang, Genay Pilarowski, Emilio Merlo Pich, Atsushi Nakatani, John Dunlop, Rina Baba, Satoru Matsuda, Masaki Daini, Yasushi Hattori, Shigemitsu Matsumoto, Mitsuhiro Ito, Haruhide Kimura, Hans Tomas Bjornsson

## Abstract

Kabuki syndrome (KS) is a rare cause of intellectual disability primarily caused by loss of function mutations in lysine-specific methyltransferase 2D (*KMT2D*), which normally adds methyl marks to lysine 4 on histone 3. Previous studies have shown that a mouse model of KS (*Kmt2d*^*+/βGeo*^) demonstrates disruption of adult neurogenesis and hippocampal memory. Proof-of-principle studies have shown postnatal rescue of neurological dysfunction following treatments that promote chromatin opening, however, these strategies are non-specific and do not directly address the primary defect of histone methylation. Since lysine-specific demethylase 1A (LSD1/KDM1A) normally removes the H3K4 methyl marks added by KMT2D, we hypothesize that inhibition of KDM1A demethylase activity may ameliorate molecular and phenotypic defects stemming from KMT2D loss. To test this hypothesis, we evaluated a recently developed KDM1A inhibitor (TAK-418) in *Kmt2d*^*+/βGeo*^ mice. We find that orally administered TAK-418 increases the numbers of newly born Doublecortin (DCX)^+^ cells and processes in hippocampus in a dose dependent manner. We also observe TAK-418-dependent rescue of histone modification defects in hippocampus both by Western blot and ChIP-Seq. Treatment rescues gene expression abnormalities including those of immediate early genes such as FBJ osteosarcoma oncogene (*Fos)* and FBJ osteosarcoma oncogene homolog B (*Fosb)*. After 2 weeks of TAK-418, *Kmt2d*^*+/βGeo*^ mice demonstrate normalization of hippocampal memory defects. In summary, our data suggest that KDM1A inhibition is a plausible treatment strategy for KS and support the hypothesis that the epigenetic dysregulation secondary to KMT2D dysfunction plays a major role in the postnatal neurological disease phenotype in KS.

## Introduction

Kabuki syndrome (KS) is a rare Mendelian disorder that affects multiple systems including neuro, immune, auditory and cardiac systems^1^. It is characterized by distinctive facial features, growth retardation, and mild to moderate intellectual disability^2^. Human genetics studies have revealed that this autosomal dominant/X-linked condition is caused by heterozygous/hemizygous loss of function in either of two genes with complementary function: *KMT2D* on human chromosome 12 or lysine-specific demethylase 6A (*KDM6A*) on human X chromosome^3,4^. Both of these disease genes encode histone modifiers that contribute to the opening of chromatin. The majority (>70%) of molecularly confirmed cases of KS have loss-of-function variants in *KMT2D. KMT2D* catalyzes the addition of methyl groups to lysine 4 of histone 3 (H3K4me1 and 3)^5,6^, which are marks associated with open chromatin. *KDM6A* also participates in chromatin opening by removing H3K27me3, a closed chromatin mark^7^. It is, therefore, likely that the observed gene dosage sensitivity in KS results from an imbalance between open and closed chromatin states at or around critical target genes. We hypothesize that it may be possible to restore this balance with drugs that promote open chromatin states^8^.

Our laboratory characterized a novel mouse model of KS1 (*Kmt2d*^*+/βGeo*^) and found that these mice have many features that overlap with KS patient phenotypes including craniofacial abnormalities, growth retardation and immune dysregulation^9-11^. These mice also demonstrate an ongoing deficiency of adult neurogenesis in the subgranular zone (SGZ) of the dentate gyrus (DG) in the hippocampus and hippocampal memory defects. We have previously shown that these deficiencies improve after a short term (2-3 weeks) oral treatment with a histone deacetylase inhibitor (HDACi), AR42^9^. Modulation of epigenetic modifications through dietary elevation of beta-hydroxybutyrate, an endogenous HDACi, may also rescue these same phenotypes^10^. However, both of these strategies are indirect, affecting chromatin opening through modulation of histone acetylation rather than histone methylation. Furthermore, HDACi have been poorly tolerated in clinical trials^12^ and a ketogenic diet, is very stringent and hard to implement in children that often have baseline feeding problems and growth retardation as is the case in Kabuki syndrome. Recently, genetic disruption of KDM1A, the factor that normally removes H3K4me1-2 was shown to rescue disrupted chromatin states in KMT2D-deficient embryonic stem cells^13^. Here, we use a novel specific inhibitor of KDM1A activity, TAK-418, to demonstrate *in vivo* rescue of adult DG neurogenesis, improvement of hippocampal memory deficits, chromatin remodeling, and gene expression abnormalities in *Kmt2d*^*+/βGeo*^ mice. TAK-418, also called 5-{(1*R*,2*R*)-2-[(cyclopropylmethyl)-amino]cyclopropyl}-*N*-(tetrahydro-2*H*-pyran-4-yl)thiophene-3-carboxamide monohydrochloride, was successfully progressed in preclinical development and was recently given orphan designation by the European Medical Agency (EMA) and the US Food and Drug Administration (FDA) for the treatment of KS. TAK-418 has already undergone Phase 1 studies on healthy volunteers to establish safety and tolerability (clinicaltrials.gov, NTC03228433 and NCT03501069). Therefore, the TAK418-dependent rescue of disease phenotypes in *Kmt2d*^*+/βGeo*^ mice provides support for a possible therapeutic role in KS.

## Methods

### Animals

Our mouse model, *Kmt2d*^*+/βGeo*^, also named *Mll2Gt*^*(RRt024)Byg*^, was originally acquired from Bay Genomics (University of California) but backcrossed in the Bjornsson laboratory. All experimental mice were on a fully backcrossed C57BL/6J background (99% verified using a mouse SNP chip). For treatment with TAK-418, mice were orally gavaged daily with drug (TAK-418, Takeda) solubilized in vehicle (methylcellulose) or with vehicle alone. Both drug and vehicle (methylcellulose) were shipped from Takeda. Drug was administered for 14 days for adult neurogenesis studies, after which mice were sacrificed on day 15. For behavioral studies, Morris water maze testing was initiated at day 15 after treatment start at ∼8 weeks of age, and the drug was continued throughout the behavioral studies (until at least day 23). A dose curve was initially performed with three doses (0.5 mg/kg/day, 1 mg/kg/day, 2 mg/kg/day). However, after that initial study, all other experiments were done with a dose of 1 mg/kg/day. For quantification of spleen size, evaluation was performed after 8 weeks of TAK-418 started at age of ∼8 weeks old, since splenomegaly is not observed until 12-16 weeks. For this reason, this cohort was kept on TAK-418 for ∼8 weeks for this reason and to evaluate for side effects. Mouse numbers for individual studies: ChIP-Seq/RNA-Seq: 3-4 per group; immunohistofluorescence: n = 11-12 per group; RT-qPCR: 5-6 per group; behavioral testing: 24-30 per group. Genotyping was performed using the following primers: Mll2_exon50F: CTGTGTGGAACCGCATCATTG; Mll2_exon51R: CGGTTCTGATCTGGCACAGCC; β-GeoR1: CTCAGTGCAGTGCAGTCAGG. The Mll2_exon50F and Mll2_exon51R pair amplify sequences from the wild allele of KMT2D, the Mll2_exon50F and β-GeoR1 pair amplify sequences specific for the targeted allele. All experiments were performed using mouse protocols approved by the Animal Care and Use Committee of Johns Hopkins University School of Medicine. The mouse protocols used for this study are in accordance with the guidelines used by the National Institutes of Health (NIH) for mouse care and handling.

### Perfusion and cryosectioning

Mice were sacrificed with a lethal dose of xylazine, ketamine combination, after which they were transcardially flushed with PBS (1x) with heparin and then perfused with 4% PFA/PBS. Brains were removed from the skulls and cryopreserved in 30% sucrose 0.1M phosphate solution overnight at 4°C. Brains were frozen and sectioned using a Microm HM 550 cryostat (Thermo Scientific). Sectioning was performed at 30μm intervals and every section of the brain was collected onto the slide in a series of 6 slides with 12 slices on each slide and stored in a -80°C freezer prior to use.

### Immunofluorescence staining

Pre-mounted slides were thawed and washed once in TBS (1x). Slices were briefly permeabilized in TBS(1x) with 0.4% Triton X-100 (30min), followed by blocking with TBS with 3% donkey serum and 0.05% Triton X-100 for 1hr at room temperature. Next each slide was incubated with primary antibody (in TBS with 3% donkey serum and 0.05% Triton X-100) overnight (O/N) at 4°C. After 3-5 washes in TBS with 0.05% Triton X-100, Alexa Fluor conjugated secondary antibodies with DAPI counterstain was added to the slide for 1 hour (hr) at room temperature. Slides were mounted with ProLong antifade (ThermoFisher Scientific, catalog# P36930) after several washes in TBS with 0.05% Triton X-100 and a final wash of TBS. Images were taken with LSM780 confocal microscope. Antibodies used included anti-DCX (Santa Cruz, catalog #sc-8066) and anti-c-fos (Abcam, catalog #190289).

### Granule cell layer and quantification of DCX^+^ cells

The area of the granule cell layer and DCX^+^ cells was measured using ImageJ as previously described^9^. Briefly, we traced the granule cell layer by hand and counted DCX^+^ positive cells within this area in all slides. Counting was performed by an investigator that was blinded to genotype and/or drug exposure.

### Western blotting

Hippocampus was dissected, snap frozen and kept in -80°C. Histones were extracted following a published acid extraction protocol^14^. Briefly, 250μl of TED buffer (0.5% Triton x 100, 2mM PMSF and 0.02% NaN3 in PBS) was added and hippocampal tissue was disaggregated by 50-60 strokes with a pellet pestle. Nuclear pellets were collected at 10,000rpm for 1 minute (min) at 4°C and resuspended in 50μl of acid extraction buffer (0.5N HCl in 10% glycerol). After O/N incubation in acid extraction buffer, supernatant was collected and histones were precipitated with acetone O/N at -20°C. Histone pellets were subsequently dissolved in water. For whole cell lysate, we disaggregated hippocampal tissue in RIPA buffer (150mM NaCl, 1.0% NP-40, 0.5% Na Deoxycholate, 0.1% SDS, 50mM Tris, pH8.0) by 50-60 strokes with pellet pestle. After incubation for 2h at 4°C, supernatant was collected. Amount of protein was quantified using a BSA protein assay. Proteins were fractionated in 4-12% NuPAGE bis-tris gel. Antibodies used included the following: anti-histone H3 (Cell Signaling Technologies, catalog #3638), anti-H3K4me3 (Cell Signaling Technologies, catalog # 9727), anti-H3K4me2 (Cell Signaling Technologies, catalog #9725), anti-H3K4me1(Abcam, catalog #ab8895), anti-c-fos (Abcam, catalog #190289), anti-phospho-p44/42 MAPK(Erk1/2) (Cell Signaling Technologies, catalog #9106), anti-p44/42 MAPK(Erk1/2) (Cell Signaling Technologies, catalog #9102), anti-β-Actin (Cell Signaling Technologies, catalog#3700).

### Behavioral testing

No exclusion criteria were used other than decreased visual acuity as evaluated by visible platform training (see below). Investigator was blinded to genotype and drug exposure status during testing. Morris water maze testing: Mice were placed in a 1.1 meter diameter tank filled with room temperature water dyed with nontoxic white paint. For analytical purposes, the tank was divided into four quadrants, with one quadrant containing a small platform submerged 1.5 cm beneath the water. On each day of training, mice were placed in the tank in a random quadrant facing away from the center and allowed to swim until they found the platform. Once they reached the platform they were left there for 30 seconds (s). If they did not reach the platform after 60s, they were placed on it for 30s. Each mouse was given four trials per day (for 5 days) with an inter-trial interval of 5-20 m and subsequently returned to its home cage. Latency to reach the platform was measured during each trial. The day after the final day of training, the platform was removed for a probe trial where mice were placed in the tank for 90s. The average number of crossings of the platform’s previous location was recorded. Visible/flagged platform training was also performed for 3 days right before the hidden platform to ensure no problems with sight. During visible training a visible flag was placed on the submerged platform, and the time for each mouse to reach the platform was measured during each 60-s trial, four of which were run in the same way as the hidden platform training. For all training and probe testing, data were recorded both manually and electronically with ANY-maze software (San Diego Instruments) when applicable. Differences in the number of platform crossings, the latency of the first crossing of the platform during the probe trial were compared between groups with a Student’s t test with significance value set at p < 0.05.

### RNA-Seq/ChIP-Seq

For RNA-Seq and ChIP-Seq, 2-week old mice were given TAK-418 at a dose of 1mg/kg body weight for a period of 2 weeks. Hippocampi were harvested at the end of the 2 week treatment. RNA was extracted with Direct-zol RNA microprep kit (ZYMO Research), RNA-Seq libraries were constructed with NEBNext Ultra RNA library prep kit for Illumina. Pooled RNA libraries were sequenced on a Hiseq2500 using 150 bp paired-end sequencing. For ChIP-Seq library construction, nuclear lysate preparation and chromatin IP were followed the ENCODE protocol from Bing Ren lab with chip graded anti-H3K4me3 (Abcam, catalog #ab8580) and anti-H3K4me1 (Abcam, catalog #ab8895). After reverse cross-linking and purifying the DNA fragments, library construction was performed with the NEBNextUltraII DNA library prep kit for Illumina based on manufacturer’s recommendations. Pooled ChIP DNA libraries were sequenced on a Hiseq2500 using a flow cell for 150 bp paired-end size.

### RNA-Seq/ChIP-Seq analysis

<u>RNA-Seq</u>: Transcriptomic data collected by RNA sequencing (RNA-Seq) was analyzed to determine the genes that are present in each sample and condition, their expression levels, and the differences between expression levels among different experimental conditions. Following quality checking with the software Fastqc, reads were mapped to the mouse genome version mm10 with the alignment tool Tophat2 v.2.1.0^15^, which allows for large ‘gaps’ in the alignment, representing introns. The aligned reads were assembled with CLASS2 v.2.1.7^16^ to create partial genes and transcript models (transfrags). Transfrags from all samples were further merged with Cuffmerge and mapped to the GENCODE v.M17 gene models^17^, to create a unified set of gene annotations for differential analyses. Lastly, gene (transcript) expression levels were computed, and differentially expressed genes (transcripts) were determined separately with the tools Cuffdiff2 v.2.2.1^18^ and DESeq^19^. Differentially expressed genes were further analyzed and graphed with R, a public free resource for statistical computing and graphing. <u>ChIP-Seq</u>: The reads collected by H3K4me1 and H3K4me3 ChIP sequencing were first trimmed 5bp at 5’end with the software seqtk (https://github.com/lh3/seqtk), followed by mapping with the short sequence alignment tool bowtie2^20^. The aligned sequences were indexed with software samtools for easy viewing in IGV (Integrated Genome Viewer). The THOR software^21^ was then used to detect and analyze differential peaks in two sets of ChIP-Seq data from distinct biological conditions with replicates. HOMER^22^ and GO analysis (http://geneontology.org/) was used to annotate the peaks, and custom scripts were written to filter THOR output, gather statistics and reformat files. Differential peaks were further analyzed and graphed with R.

## Results

### Kmt2d^+/βGeo^ mice demonstrate a global decrease of histone H3K4 methylation in the hippocampus

*Kmt2d*^*+/βGeo*^ mice contain an expression cassette encoding a β-galactosidase neomycin resistance fusion protein (β-Geo) inserted into intron 50 of the *Kmt2d* gene locus on mouse chromosome 15. This cassette contains a 5’ end splice acceptor sequence and a 3’ end cleavage and polyadenylation signal. The mutated allele, therefore, results in a truncated KMT2D protein with the peptide encoded by the first 50 exons of KMT2D fused to β-Geo at the C-terminus^9^. The truncated KMT2D fusion protein, therefore, lacks the C-terminal catalytic SET domain of KMT2D, which is responsible for its H3K4 methyltransferase activity. Because KMT2D is a prominent mammalian H3K4 methyltransferase and *Kmt2d*^*+/βGeo*^ mice have disrupted hippocampal neurogenesis, we hypothesized we would see diminished levels of mono- and di-methylated H3K4 (H3K4me1/2) in the granule cell layer of the dentate gyrus in *Kmt2d*^*+/βGeo*^ mice. To test this, we performed Western blots of histone extracts from mouse hippocampi and observed significantly (p < 0.0001) reduced levels of H3K4me1 and H3K4me2 in *Kmt2d*^*+/βGeo*^ mice compared to *Kmt2d*^*+/+*^ littermates when normalized to total histone 3 (H3) (Fig. 1a-b). Therefore, although KMT2D and Lysine Methyltransferase 2C (KMT2C) have some overlapping function, our results indicate that the latter is unable to compensate for the heterozygous loss of *Kmt2d* in the hippocampus.

**Fig. 1:**
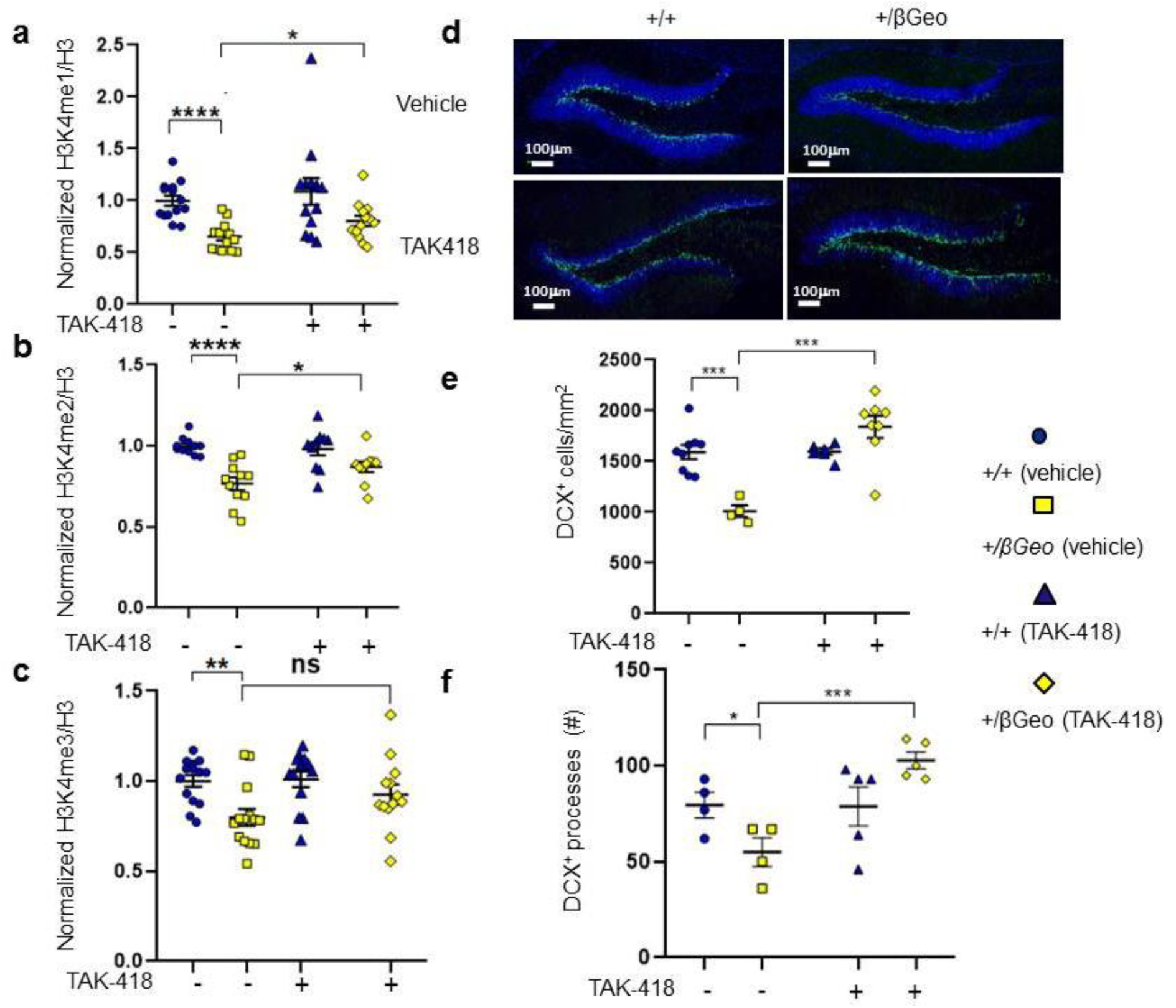
TAK-418 rescues histone H3K4 methylation abnormalities and neurogenesis defects in hippocampus of *Kmt2d^+/βGeo^* mice. Quantification of H3K4me1, H3K4me2, and H3K4me3 levels from Western blots of hippocampal lysates from both genotypes with and without 2 weeks of TAK-418 treatment (**a-c**). A deficiency of DCX^+^ cells in the granule cell layer of the hippocampus seen in *Kmt2d*^*+/βGeo*^ mice compared to littermates normalizes after 2 weeks of TAK-418 (**d, e**). A defect of the number of DCX^+^ processes is also rescued after 2 weeks on TAK-418 (**f**). Student’s t-test. *p < 0.05,**p < 0.01, ***p < 0.005, ****p < 0.001.

Previously it has been reported that KMT2D is required for H3K4 trimethylation (H3K4me3) in bivalent promoters^23^. Indeed, we find that the level of H3K4me3 is also reduced (Fig. 1c), and that the ratio of H3K4me3 to total histone 3 is significantly reduced in *Kmt2d*^*+/βGeo*^ mice compared to *Kmt2d*^*+/+*^ littermates (p < 0.0016), although to a lesser extent than the observed deficiencies of H3K4me1/2.

### TAK-418 treatment rescues the deficiency of mono- and di-methylated, but not tri-methylated, H3K4 in the hippocampus of *Kmt2d*^*+/βGeo*^ mice

TAK-418 is a selective inhibitor of the lysine-specific demethylase 1(KDM1A/LSD1) recently developed by Takeda Pharmaceutical Company Limited. KDM1A, in association with the corepressor of REST1 (CoREST) complex, removes methylation at H3K4me1/2 sites^24-27^. Previously, genetic disruption of *KDM1A* was found to rescue differentiation defects in KMT2D-deficient embryonic stem cells^13^. To test whether TAK-418 is able to rescue epigenetic abnormalities *in vivo* in the hippocampus, mice were given TAK-418 at 1mg/kg/day by oral administration for two weeks. Histones were then extracted from dissected hippocampus for analysis. We observed significant rescue (p < 0.05) by TAK-418 treatment of the deficient H3K4me1 levels in *Kmt2d*^*+/βGeo*^ mice compared to *Kmt2d*^*+/+*^ littermates (Fig. 1a). Similarly, we observed a significant rescue (p < 0.05) of H3K4me2 levels in *Kmt2d*^*+/βGeo*^ mice with TAK-418 compared to littermates on vehicle (Fig. 1b). H3K4me3 levels were also increased by TAK-418, albeit not to statistical significance (Fig. 1c). Thus, in summary, we observe biochemical rescue of H3K4 methylation in hippocampus, a disease relevant tissue. As expected, effects of TAK-418 on all three H3K4 methylation marks were largely correlated.

### TAK-418 rescues defects of adult neurogenesis in *Kmt2d*^*+/βGeo*^ mice

Adult hippocampal neurogenesis is an ongoing process that persists throughout life^28,29^ in which adult-born neural progenitors in the subgranular zone (SGZ) give rise to excitatory granule cell neurons residing in the hippocampal dentate gyrus (DG)^29^. The neuroblast migration protein DCX is highly expressed in immature neurons and sharply decreases with maturation of neurons^30,31^, providing a cell stage/type-specific marker to quantify adult neurogenesis *in vivo*. We tested the effects of TAK-418 on adult neurogenesis as measured by DCX^+^ cells per mm^2^ in the DG and SGZ. As before^9,10^, we find decreased numbers of DCX^+^ cells in *Kmt2d*^*+/βGeo*^ mice compared to *Kmt2d*^*+/+*^ littermates. On TAK-418 treatment, this phenotype showed dose-dependent normalization (Supplementary Fig. 1) with full rescue at the medium dose (1 mg/kg/day, Fig. 1d, e). Dendrites can be visualized in some DCX^+^ cells, as DCX is a cytoplasmic protein that is associated with microtubules present in dendrites^32,33^. We counted the DCX^+^ cells with dendrites and found that *Kmt2d*^*+/βGeo*^ mice have fewer DCX^+^ cells with dendrites compared to *Kmt2d*^*+/+*^ littermates (Fig. 1f), although this likely, to some extent, reflects absolute cell numbers. *Kmt2d*^*+/βGeo*^ mice treated with TAK-418 have significantly increased numbers of DCX^+^ cells with dendrites compared to *Kmt2d*^*+/βGeo*^ mice treated with vehicle (Fig. 1f). The dendrites in the *Kmt2d*^*+/βGeo*^ mice also appeared shorter than the dendrites in *Kmt2d*^*+/+*^ littermates, which reached farther into the granule cell layer and even into the molecular cell layer. TAK-418 treatment restored the dendrite length and characteristics in *Kmt2d*^*+/βGeo*^ mice resulting in longer dendrites extending throughout the granule cell layer and into the molecular cell layer (Supplementary Fig. 2).

### TAK-418 treatment rescues the genome-wide deficiency of H3K4 methylation in *Kmt2d*^*+/βGeo*^ mice

Given the global H3K4 methylation deficiency and rescue by TAK-418 in the hippocampus of *Kmt2d*^*+/βGeo*^ mice, we next interrogated the genome-wide histone profiles in hippocampus by chromatin immunoprecipitation and high-throughput sequencing (ChIP-Seq). We first performed H3K4me1-ChIP-Seq on hippocampi harvested from *Kmt2d*^*+/βGeo*^ and *Kmt2d*^*+/+*^ mice treated with either vehicle or TAK-418. The majority of H3K4me1 peaks (∼57.7%) appeared in intergenic regions which is consistent with known enhancer-associated roles of this mark^5^. A third of peaks (∼31.2%) appeared to be in introns, and fewer were at the promoter/transcription start site (∼2.7%) or transcription start site (∼1.8%). There was no obvious difference between the two genotypes. In untreated hippocampi, we observed a total of 621 differentially bound peaks of H3K4me1 in *Kmt2d*^*+/βGeo*^ mice relative to *Kmt2d*^*+/+*^ (Fig. 2a, Supplementary Table S1). Of these, 396 peaks were decreased and 225 were increased in vehicle-treated *Kmt2d*^*+/βGeo*^ compared to *Kmt2d*^*+/+*^ littermates (Fig. 2a). The 28 most statistically significant peaks are shown in Supplementary Table S1. In general, *Kmt2d*^*+/+*^ mice had more differentially H3K4me1-bound peaks compared to *Kmt2d*^*+/βGeo*^ mice when either genotype was on vehicle (396/225, 1.76/1, Fig. 2a). Upon treatment, we observed 622 differentially bound H3K4me1 peaks, with 322 increased and 300 decreased in *Kmt2d*^*+/βGeo*^ on TAK-418 compared to *Kmt2d*^*+/+*^ mice on vehicle (Fig. 2b, Supplementary Table S2). The ratio of bound H3K4me1 in *Kmt2d*^*+/βGeo*^ on TAK-418 compared to *Kmt2d*^*+/+*^ mice on vehicle was 1 to 0.93 (Fig. 2b). An inverse linear regression revealed a correlation coefficient of -0.632 (Fig. 2c) of log2 fold change of the common bound loci between the comparison of *Kmt2d*^*+/+*^*/Kmt2d*^*+/βGeo*^ on vehicle and *Kmt2d*^*+/βGeo*^/*Kmt2d*^*+/βGeo*^ on TAK-418. These results indicate that TAK-418 treatment leads to generalized, albeit partial, rescue of the H3K4me1 deficiency.

**Fig. 2:**
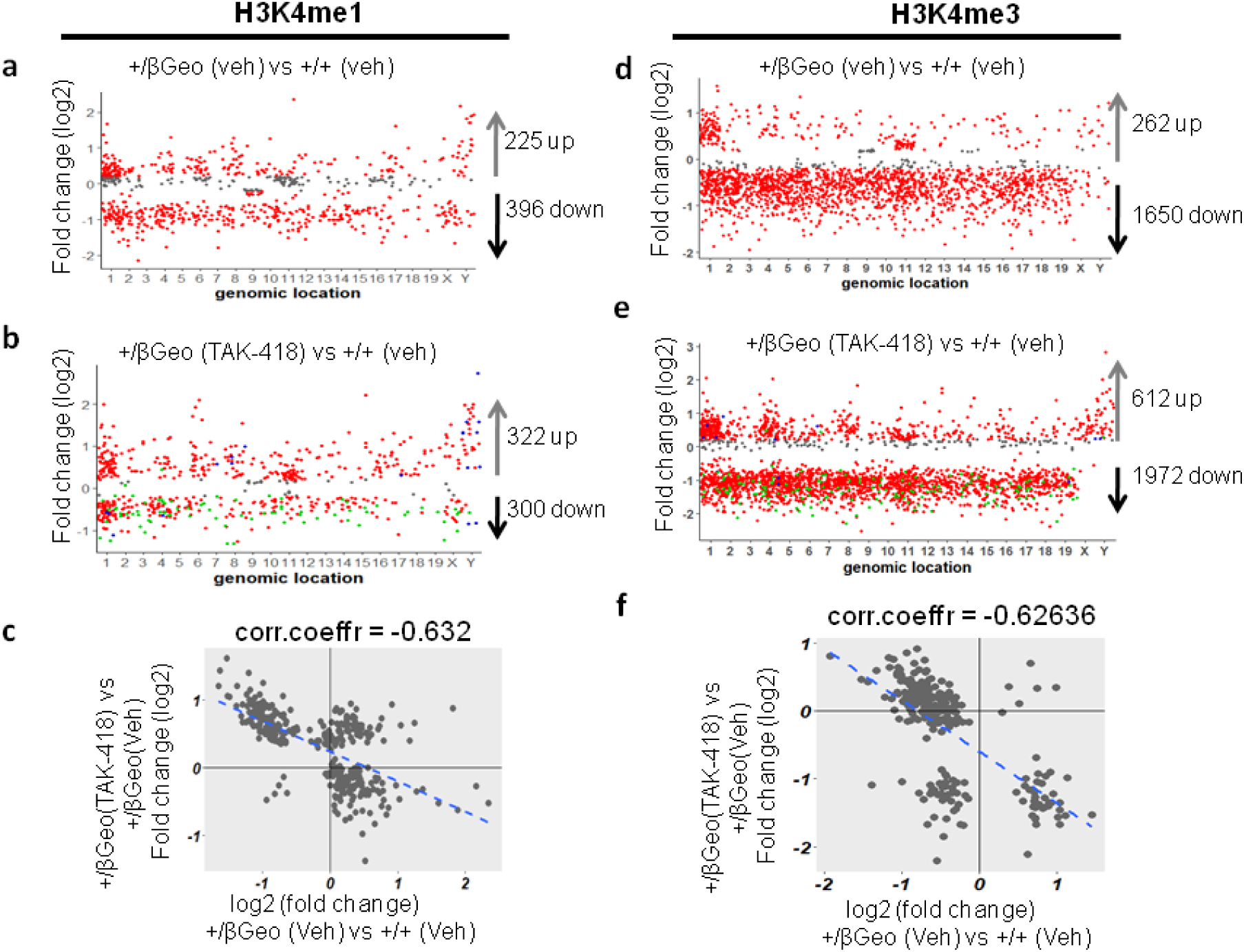
Hippocampal genome wide levels of H3K4me1 and H3K4me3 demonstrate TAK-418-dependent rescue. Comparison of hippocampal H3K4me1 levels of *Kmt2d*^*+/βGeo*^ on vehicle or TAK-418 compared to *Kmt2d*^*+/+*^ on vehicle (**a, b, c**). Comparison of hippocampal H3K4me3 levels of *Kmt2d*^*+/βGeo*^ on vehicle or TAK-418 compared to *Kmt2d*^*+/+*^ on vehicle (**d, e, f**). Each point corresponds to a genomic location of a peak with a statistically significant difference between the two genotypes. Log2 fold change >0.2 or <0.2 are in red, and others are in gray. To allow visualization of changes upon TAK-418 treatment the green dots indicate H3K4me1/3-bound locations that were significantly increased in *Kmt2d*^*+/βGeo*^ mice on vehicle (**b**,**e**) and blue dots indicate H3K4me1/3-bound peaks that were decreased in *Kmt2d*^*+/βGeo*^ on vehicle (**b**,**e**).

We next performed H3K4me3 ChIP-Seq. As expected the majority of H3K4me3 peaks are intragenic, with ∼18-19% in the promoter region, ∼16-17% peaks in exons, ∼33-35% peaks in CpGs in introns and ∼7% intergenic. There was no obvious difference in overall distribution between the two genotypes. We observed 262 loci that were increased and 1650 loci that were decreased in the *Kmt2d*^*+/βGeo*^ mice compared to the *Kmt2d*^*+/+*^ mice when vehicle treated (Fig. 2d, Supplementary Table S3). On average there is much more H3K4me3 bound in the *Kmt2d*^*+/+*^ mice compared to *Kmt2d*^*+/βGeo*^ mice with a ratio of 6.63/1(Fig. 2d, both genotypes on vehicle). In contrast, when we compared *Kmt2d*^*+/βGeo*^ treated with TAK-418 to *Kmt2d*^*+/+*^ treated with vehicle control, we observed 612 significantly increased loci and 1972 significantly decreased loci in *Kmt2d*^*+/βGeo*^ compared to *Kmt2d*^*+/+*^ littermates. As before, we observe more H3K4me3 binding upon treatment (3.22/1, *Kmt2d*^*+/βGeo*^ on TAK-418/ *Kmt2d*^*+/+*^ on vehicle, Fig. 2e, Supplementary Table S4). An inverse linear regression revealed a correlation coefficient (r) of -0.62636 upon plotting log2 fold change of the common bound loci between the comparison of *Kmt2d*^*+/+*^/*Kmt2d*^*+/βGeo*^ on vehicle and *Kmt2d*^*+/βGeo*^/*Kmt2d*^*+/βGeo*^ on TAK-418, suggesting partial rescue of the genome-wide-deficiency of H3K4me3 by TAK-418 (Fig. 2f).

### Global gene expression changes are rescued in *Kmt2d*^*+/βGeo*^ mice on TAK-418

We next interrogated functional effects of the observed differential histone modifications, as measured by changes in gene expression, in order to define a list of potential KMT2D target genes in the hippocampus. We performed RNA-Seq on samples from whole hippocampi harvested from *Kmt2d*^*+/βGeo*^ mice and *Kmt2d*^*+/+*^ littermates treated with or without TAK-418. Differential gene expression analysis between *Kmt2d*^*+/βGeo*^ mice and *Kmt2d*^*+/+*^ littermates treated with vehicle control revealed a total of 136 differentially expressed genes (DEGs) (absolute log2 fold change >0.5, Fig. 3a, Supplementary Table S5). Among the 136 DEGs, 74 genes were downregulated and 62 genes were upregulated in hippocampus from *Kmt2d*^*+/βGeo*^ mice compared to *Kmt2d*^*+/+*^ littermates (Fig. 3a). Genes that were downregulated in *Kmt2d*^*+/βGeo*^ mice compared to *Kmt2d*^*+/+*^ littermates were enriched for networks of ionic transport and negative regulation of synaptic signaling (Fig. 3b). The pathways affected by genes upregulated in *Kmt2d*^*+/βGeo*^ mice compared to *Kmt2d*^*+/+*^ littermates included tissue development, adhesion and extracellular structure organization (Fig. 3b). After treatment with TAK-418, however, we observed a reversed pattern of expression effects. Among 130 DEGs, only 38 genes were downregulated and 92 genes were upregulated in *Kmt2d*^*+/βGeo*^ mice on TAK-418 compared to *Kmt2d*^*+/+*^ littermates treated with vehicle control (Fig. 3c, Supplementary Table S6). Pathways enriched among genes upregulated in *Kmt2d*^*+/βGeo*^ mice treated with TAK-418 were involved in neurogenesis and neuronal projection development (Fig. 3d). The majority of DEGs (absolute log2 fold change <0.5) in *Kmt2d*^*+/βGeo*^ mice compared to *Kmt2d*^*+/+*^ littermates were normalized with TAK-418 treatment (blue dots, Fig. 3c). Among the genes that were highly differentially expressed in untreated *Kmt2d*^*+/βGeo*^ animals and rescued with TAK-418 were the immediate early genes *Fos* and *Fosb*, in addition to Neurexophilin-3 (*Nxph3)*, relaxin/insulin-like family peptide receptor 1 *(Rxfp1)*, down-stream targets of extracellular-signal-regulated kinase (ERK) signaling, and ribosomal protein genes (Fig. 3e). When we plotted the log2 fold change of the common genes found in the comparison of *Kmt2d*^*+/βGeo*^*/Kmt2d*^*+/+*^ on vehicle and *Kmt2d*^*+/βGeo*^ on TAK-418/*Kmt2d*^*+/βGeo*^ on vehicle, we observe a highly significant inverse correlation (R = -0.9) indicating that TAK-418 treatment of *Kmt2d*^*+/βGeo*^ mice rescued the disrupted expression levels of genes in untreated *Kmt2d*^*+/βGeo*^ mice to approximately what is normally seen in *Kmt2d*^*+/+*^ littermates (Fig. 3f).

**Fig. 3:**
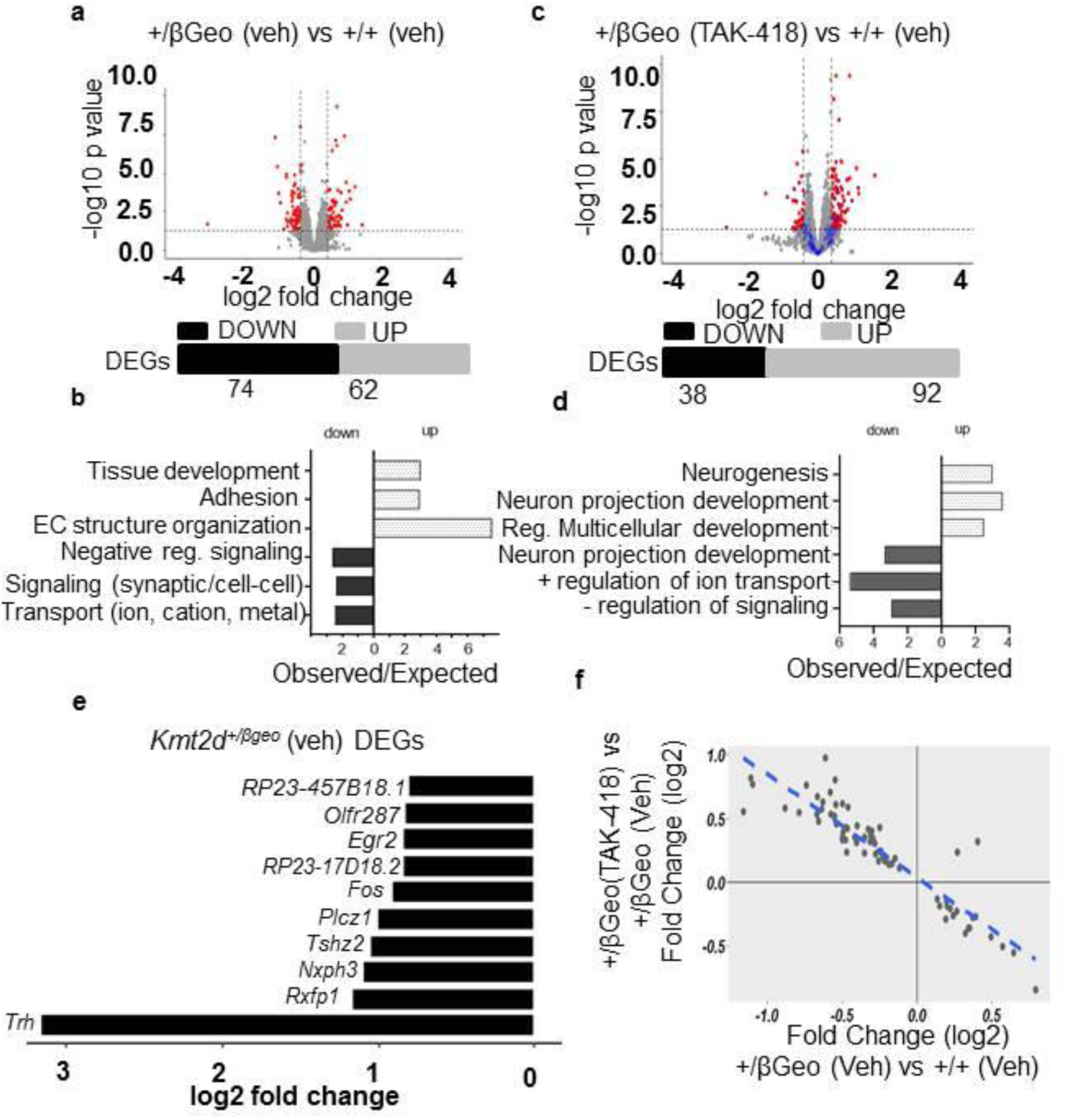
TAK-418 rescues gene expression abnormalities in *Kmt2d^+/βGeo^* mice. An overview of gene expression abnormalities (volcano plot, **a**) and a summary of major gene categories (**b**) among differentially expressed genes (DEGs) in hippocampus when comparing *Kmt2d*^*+/βGeo*^ and *Kmt2d*^*+/+*^ on vehicle. Many of the DEGs from vehicle treatment (**a**) were normalized (blue dots in c) upon a comparison of hippocampal gene expression levels between *Kmt2d*^*+/βGeo*^ on TAK-418 and *Kmt2d*^*+/+*^ on vehicle (**c, d**). Red dots are differentially expressed genes with absolute log2 fold change≥0.5 and P-value<0.05. A view of representative (top 10, all downregulated) DEGs between *Kmt2d*^*+/βGeo*^ and *Kmt2d*^*+/+*^ on vehicle (**e**). A correlation of DEGs in *Kmt2d*^*+/βGeo*^ (TAK-418/vehicle) and the two genotypes (*Kmt2d*^*+/βGeo*^/*Kmt2d*^*+/+*^ on vehicle) reveals TAK-418-dependent rescue (**f**).

### The immediate early gene *Fos* shows decreased expression in *Kmt2d*^*+/βGeo*^ mice that is rescued with TAK-418

We validated gene expression of some of the most interesting candidate genes by RT-qPCR. NXPH3 is a specific ligand of synaptic alpha-neurexins and is essential for efficient neurotransmitter release^34^. *Nxph3* expression is significantly downregulated (p < 0.009) in *Kmt2d*^*+/βGeo*^ mice compared to *Kmt2d*^*+/+*^ littermates (Fig. 4a). Upon treatment with TAK-418, we saw a modest increase of *Nxph3* levels (p < 0.09, Fig. 4a). *Fos* and *FosB* are immediately early genes that respond to extracellular stimuli. *Fos*, a marker of neuronal activity, has been associated with a number of neural and behavioral responses to acute stimuli expression. *Fos* expression is downregulated in *Kmt2d*^*+/βGeo*^ mice (p < 0.05), however, upon treatment with TAK-418, the expression level of *Fos* is upregulated significantly (p < 0.03) (Fig. 4b). *FosB* expression trends mirrored those of *Fos*, but to a lesser degree (Fig. 4c). At the protein level, we observed a marginal decrease of FOS protein in total hippocampus lysate of *Kmt2d*^*+/βGeo*^ mice compared with *Kmt2d*^*+/+*^ littermate controls and a marginal increase of FOS protein upon treatment with TAK-418 in *Kmt2d*^*+/βGeo*^ mice (Fig. 4d, e). We hypothesize that the incomplete rescue could be related to the differential cell type composition in samples from brain tissue. In support of this hypothesis, when we stained the mouse brain slices for FOS and counted FOS^+^ positive cells and calculated the average number of FOS^+^ positive cell per mm^2^ from 9-10 slices per mouse with 3-5 mice each examination group (Supplementary Fig. 3), we noticed significantly fewer FOS^+^ positive cells per mm^2^ of dentate gyrus in *Kmt2d*^*+/βGeo*^ compared with *Kmt2d*^*+/+*^ littermate controls (Fig. 4f, g). After 2 weeks of treatment with TAK-418 in *Kmt2d*^*+/βGeo*^ mice, the average FOS^+^ positive cells per mm^2^ increased, although not to a significant level (Fig. 4f, g and Supplementary Fig. 3). *Fos* mediates a signaling cascade involving phosphorylation by extracellular signal-regulated kinases (ERKs). Activated (phosphorylated) ERK is known to be one of the regulators of *Fos* expression and activity. The decrease in *Fos* RNA and protein levels in *Kmt2d*^*+/βGeo*^ mice prompted us to look at the phospho-ERK level in total hippocampal lysate. The ratio of phosphorylated ERK to total ERK was decreased significantly in *Kmt2d*^*+/βGeo*^ mice compared with *Kmt2d*^*+/+*^littermate (Supplementary Fig. 4), supporting the hypothesis that the ERK based activation of the Fos signaling cascade is disrupted in *Kmt2d*^*+/βGeo*^ mice.

**Fig. 4:**
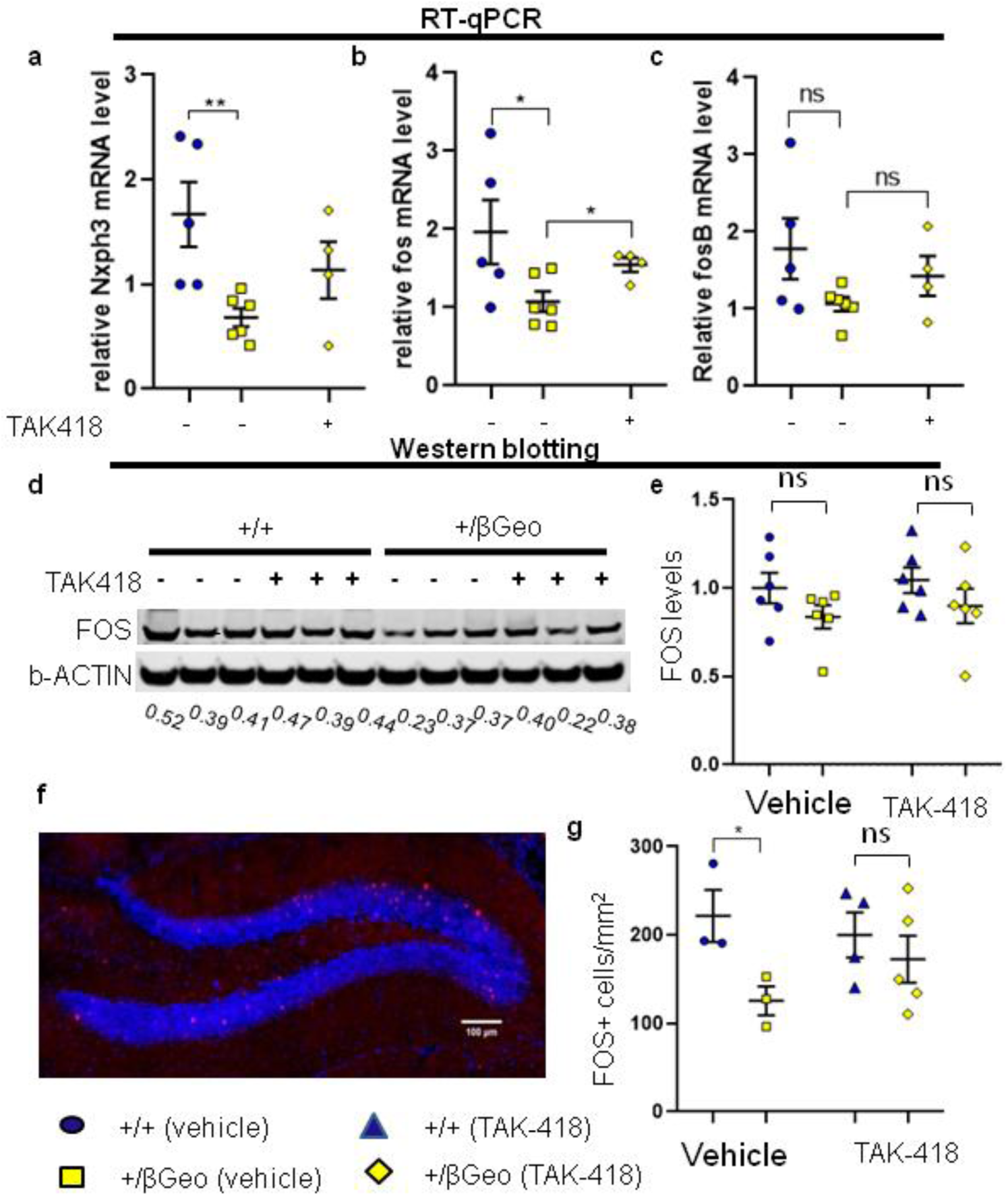
Gene expression abnormalities of disease-relevant candidate genes in *Kmt2d^+/βGeo^* mice are reflected in abnormal RT-qPCR, Western blot, and immunofluorescence staining. *Nxph3, fos* and *fosb* are disease-relevant DEGs and all demonstrate consistent decreased gene expression in *Kmt2d*^*+/βGeo*^ mice compared to littermates and these changes rescue with TAK-418 (**a, b, c**). Western blotting of FOS reveals a defect at the protein level in *Kmt2d*^*+/βGeo*^ mice compared to littermates (with some rescue, **d, e**). Immunofluorescence staining of tissue slices from *Kmt2d*^*+/βGeo*^ mice and littermates demonstrate a significant defect that rescues upon TAK-418 treatment (**f, g**). f is a representative immunofluorescence image of FOS from *Kmt2d*^*+/βGeo*^ treated with TAK-418. Student’s t-test, *p < 0.05,**p < 0.01.

### TAK-418 rescues the visuospatial learning and memory defect in *Kmt2d*^*+/βGeo*^ mice

Adult-born hippocampal neurons become integrated in the DG circuitry where they mediate neuronal plasticity in support of visuospatial learning and pattern discrimination. Previously we have shown that *Kmt2d*^*+/βGeo*^ mice have defects in spatial learning and memory formation as evaluated by a Morris water maze test^9^. Here we confirm^9,10^ that vehicle treated *Kmt2d*^*+/βGeo*^ mice persistently crossed the platform less frequently (p < 0.0007, Fig. 5a) than vehicle treated *Kmt2d*^*+/+*^ littermates in the Morris water maze. We also found that *Kmt2d*^*+/βGeo*^ mice demonstrate a longer latency to reach the platform (p < 0.0031, Fig. 5b) and spend less time on the platform (Fig. 5c) and in the quadrant with the platform (p < 0.0026, data not shown) compared to *Kmt2d*^*+/+*^ littermates. The TAK-418 treatment considerably ameliorated the defect observed in *Kmt2d*^*+/βGeo*^ mice in the Morris water maze test (Fig. 5a). In the final probe trial, the *Kmt2d*^*+/βGeo*^ mice treated with TAK-418 had increased frequency of crossing the platform compared to untreated *Kmt2d*^*+/βGeo*^ mice (p < 0.0206, Fig. 5a). Similarly, *Kmt2d*^*+/βGeo*^ treated with vehicle consistently had increased latency to find the platform (p < 0.0088, Fig. 5b) and spent less time on the platform (p < 0.0131, Fig. 5c) both of which were rescued with TAK-418. There didn’t appear to be any obvious confounders of either drug or genotype on vision or muscle strength based on the results collected from the Morris water maze. Specifically, there were no major side effects on muscles or vision after treatment with TAK-418 for 2-3 weeks, as the average swimming speed was very similar in both genotypes on and off TAK-418 (Fig. 5d) and there was no difference in the 3-day visual trial with regard to the latency to find the platform (Fig. 5e). During the five days of training, the mice improved over time demonstrating appropriate learning, however, the *Kmt2d*^*+/βGeo*^ mice appeared to perform worse (Supplementary Fig. 5) and were significantly worse on day 4 (p < 0.05) but appeared to demonstrate some rescue on TAK-418, although it was not significant (repeated measures ANOVA).

**Fig. 5:**
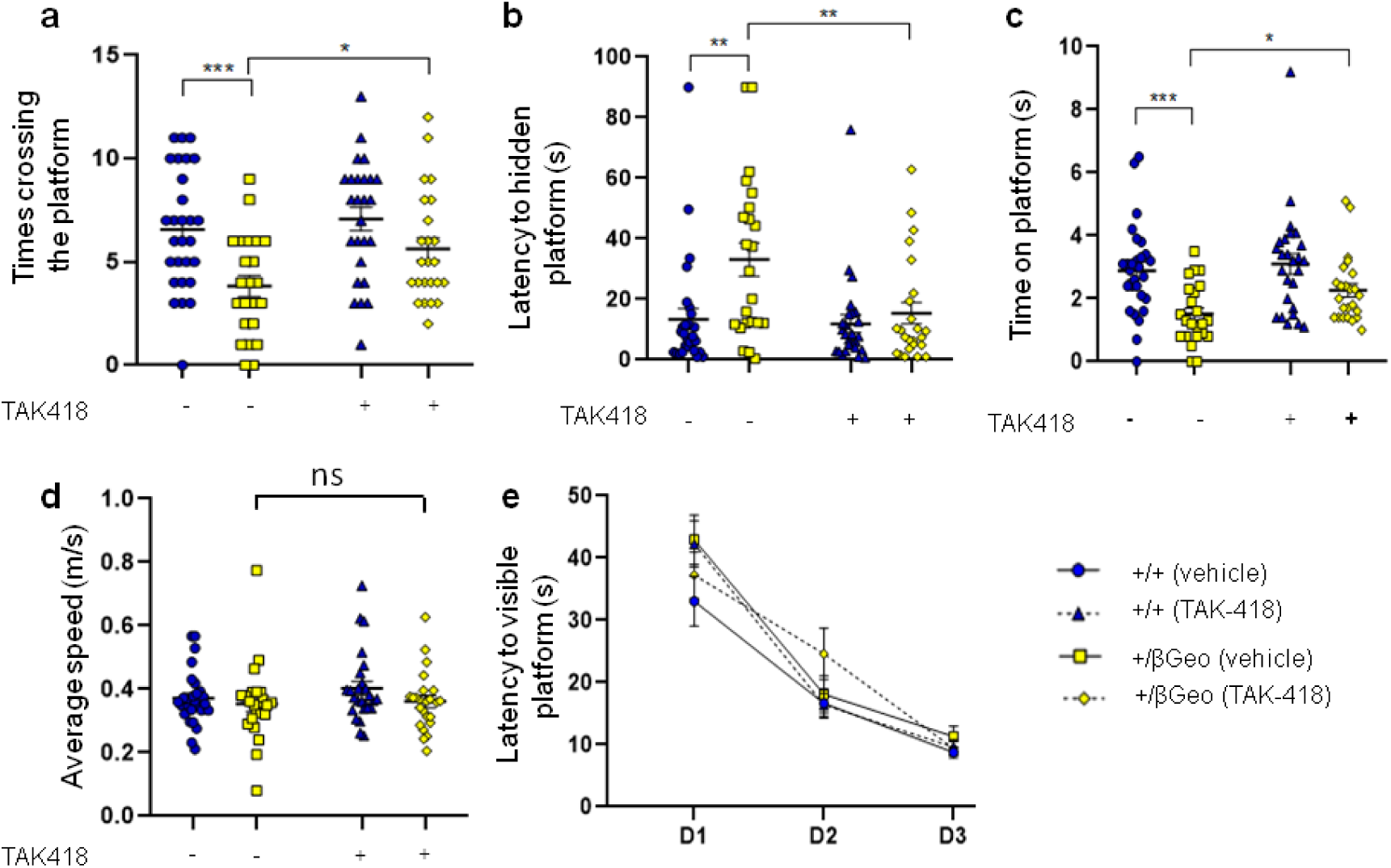
TAK-418 rescues the visual spatial learning and memory defect in *Kmt2d^+/βGeo^* mice. *Kmt2d*^*+/βGeo*^ mice have significant abnormalities in the number of times crossing platform area during probe trial (**a**), latency to find platform (**b**) and time spent on platform (**c**); all defects are rescue on TAK-418. These results are not confounded by muscle strength or vision, as *Kmt2d*^*+/βGeo*^ mice have similar average speed (**d**) and no significant defects in a visual flag finding regimen compared to *Kmt2d*^*+/βGeo*^ mice (**e**), t-test (**a-d**), Repeated measures ANOVA (**e**), *p < 0.05,**p < 0.01, ***p < 0.005.

### TAK-418 is well tolerated in mice at therapeutic levels and leads to rescue of splenomegaly, another disease relevant phenotype

We observed no obvious side effects in the mice that were on TAK-418, including no effect on weight or general well-being (data not shown). We have previously observed splenomegaly at or around 4 months of age (data not shown). Given this finding, we kept a cohort of mice on TAK-418 for 8 weeks (starting at 2 months of age) and again observed no obvious side effect or impact on weight (no weight loss or weight gain). We did, however, observe splenomegaly in the *Kmt2d*^*+/βGeo*^ mice treated with vehicle, but correction of the splenomegaly in *Kmt2d*^*+/βGeo*^ mice on TAK-418 (Supplementary Fig. 6) indicating that this phenotype may also be malleable by TAK-418 treatment.

## Discussion

Post-translational modifications of histone proteins, such as acetylation, methylation, phosphorylation, and ubiquitination are thought to serve as crucial regulatory signals to control gene expression in eukaryotic cells and are important for maintaining genomic integrity^35^. Emerging evidence suggests that dysregulation of epigenetic modifications is mechanistically linked to both cancer and developmental defects including neurodevelopmental disorders^36^. Histone methylation, in particular, confers active or repressive chromatin states in a locus-specific manner. The histone methylation state is meticulously regulated by the balance between two opposing enzyme systems: lysine methyltransferases and lysine demethylases (KMTs and KDMs). Disease-causing variants in KMTs and KDMs have been identified in multiple disorders in patients with intellectual disability, indicating that altered regulation of histone methylation can lead to intellectual disability^37^. Pharmaceutical inhibition of epigenetic targets counteracting the epigenetic effects of these loss-of-function variants has been investigated as a possible therapeutic option in several epigenetic conditions including KS^9,10,38^.

In recent years, data have emerged that support the notion that KMT2D dysfunction affects cellular function in the hippocampus. These include several studies that report more severe visuospatial disruption (linked to hippocampus) in molecularly confirmed patients with KS compared to other individuals with non-KS intellectual disability^39,40^. MRI images also suggest a grossly smaller hippocampus in individuals with molecularly confirmed KS^41^. These studies help set the stage for a potential clinical trial, as do recently developed international diagnostic criteria for KS^2^. These data also support the strategy to focus on hippocampus and dentate gyrus to estimate the effect of treatment, although they in no way exclude that other brain regions or cell types play a role in the neurological disease phenotype in KS.

Recently, genetic targeting of *KDM1A* was found to rescue a cellular phenotype observed with loss of function of *KMT2D* in embryonic stem cells^13^. KDM1A, also known as LSD1 or AOF2, is the first identified FAD-dependent histone demethylase capable of specifically demethylating mono- and di-methylated lysine 4 of histone H3 (H3K4me1 and H3K4me2) the very marks placed by KMT2D^24^. This suggests the feasibility of a more targeted therapeutic strategy for Kabuki syndrome, namely KDM1A inhibition. KDM1A associates with HDAC1/2,CoREST, BHC80, and BRAF35^25^. BRAF35, with its HMG DNA-binding domain, is thought to recruit the KDM1A/CoREST complex to the target sites. Subsequent deacetylation of the histone tail by the HDAC1/2 at the target sites then enables KDM1A to demethylate H3K4^26^. It has been demonstrated that hyperacetylated nucleosomes are less susceptible to CoREST/LSD1-mediated demethylation^42^, suggesting that hypoacetylated nucleosomes may be the preferred physiological substrates. We have previously shown that AR-42, a histone deacetylase inhibitor (HDACi), can rescue the learning and memory defect in *Kmt2d*^*+/βGeo*^ mice^9^. However, AR-42, a pan histone deacetylase inhibitor, likely impacts deacetylase activities indiscriminately across a range of distinct HDAC-containing multiprotein complexes. Such broad cellular effects may result in a narrow therapeutic window between disease efficacy and toxicity. Indeed, when we treated *Kmt2d*^*+/βGeo*^ mice with AR-42, we began to observe the effect at 5mg/kg/d with full effect obtained at 10mg/kg/d. At 25mg/kg/d, however, we started to observe cytotoxicity effect, reflected by fewer DCX^+^ positive cells compared with *Kmt2d*^*+/βGeo*^ mice treated with vehicle^9^. We had postulated that the treatment effect was likely indirect, primarily affecting histone acetylation, with a secondary effect on histone methylation^9^. An alternative hypothesis is that AR-42 exerts its effect through inhibition of HDAC1/2 activity in KDM1A-CoREST complex, leading to hyperacetylation of the target sites which are more resistant to KDM1A-mediated demethylation, thus retaining the H3K4 methylation. This may mean that low dose combined treatment with deacetylase inhibitor and demethylase inhibitor, or dual histone deacetylase and demethylase inhibitors targeting the CoREST complex may have synergistic effect and be a particularly effective treatment for Kabuki syndrome. In contrast to AR-42, TAK-418, received full effect at 1mg/kg/d with regards to DCX^+^ positive cells and no adverse effect have been observed with higher dose of TAK-418. Thus, TAK-418, as specific inhibitor to KDM1A, may also have the potential to be a single agent treatment for KS through its effects on H3K4 methylation.

In addition to the expected global increase of H3K4me1 and H3K4me2 after treatment with TAK-418 in *Kmt2d*^*+/βGeo*^, we also observed a global increase of H3K4me3, likely due to the accumulation of H3K4me2 and subsequent conversion into H3K4me3 by H3K4 methyltransferases. Alternatively, the KDM1A complex has been shown to be unstable while binding to methylated targets sites. iBRAF, the paralogue of BRAF35^43^, may compete with BRAF35 for the same target sites and recruit KMT2A complex, which subsequently may enhance trimethylation of H3K4, however, this will need to be explored in future studies. Although, KDM1A-inhibiting treatment is promising for Kabuki syndrome, the dose will need to be optimized. This is obvious because missense mutations in *KDM1A* have been identified in three individuals with developmental delay^44-46^, indicating that too much KDM1A inhibition can be damaging. The developmental symptoms in these individuals are similar to those of Kabuki syndrome (MIM: 147920), characterized by distinct craniofacial features including widely spaced teeth and palatal abnormalities, indicate that too much disruption of KDM1A actively could also be detrimental to intellectual function. Although this protein also has non-epigenetic function, it is likely that the cause of the phenotypes relates to its epigenetic function because biochemical studies have demonstrated that these mutant proteins exhibit reduced stability and demethylase activity^46^, indicating a loss-of-function mechanism.

Our RNA and ChIP sequencing data indicate that the immediate early genes (IEGs) such as *Fos* and *FosB* may be one class of genes that is affected in Kabuki syndrome and show rescue on TAK-418. Rusconi, *et al.* have shown that the KDM1A complex interacts with serum response factor (SRF) under resting conditions and an enrichment of KDM1A complex with SRF has been detected in *c-fos* promoter which contains the serum response element (SRE). SRF is known to be constitutively bound to the DNA of its target genes and the interaction of the KDM1A complex with SRF modulates the H3K4 methylation level at the *fos* promoter^47,48^. We also noted gene expression changes that would be consistent with decreased ERK signaling and, in fact, ERK activation appears deficient in *Kmt2d*^*+/βGeo*^ hippocampi compared to those in littermates. Previous data from fibroblasts from KS individuals and zebrafish models of Kabuki syndrome has implicated decreased activation of this pathway^49,50^, which is concordant with our observations. Thus, both studies reveal abnormalities of ERK signaling and future studies should further elucidate whether ERK abnormalities play a mechanistic role in the pathogenesis of Kabuki syndrome.

In summary, here we show that oral administration of a KDM1A specific inhibitor, TAK-418, can ameliorate neurological problems at the cellular, molecular, gene expression and functional levels in a mouse model of Kabuki syndrome (*Kmt2d*^*+/βGeo*^ mice). TAK-418 treatment increases adult neurogenesis in adult mice, as indicated by increased DCX^+^ cells in the granule cell layer of the hippocampus. At the molecular level, TAK-418 increases the global level of mono-, di- and tri-methylated H3K4 (H3K4me1/2/3) in *Kmt2d*^*+/βGeo*^ mice as assessed by both Western blot and ChIP-Seq. TAK-418 treatment also corrects the differential gene expression profile abnormalities found in *Kmt2d*^*+/βGeo*^ mice compared to *Kmt2d*^*+/+*^ littermates. Finally, and most importantly, we show that TAK-418 can correct the functional deficits by improving the learning and memory behavior of *Kmt2d*^*+/βGeo*^ mice. Currently we do not know if these effects will translate in humans. However, the present data are informative in respect to the TAK-418 dose range and exposure to be achieved to produce pharmacologically relevant effects in *Kmt2d*^*+/βGeo*^ mice, contributing to better design for the clinical proof of concept trial. In summary, our data support the hypothesis that KS is a treatable cause of intellectual disability and that KDM1A inhibition, may be a novel and effective mechanism of action for the treatment of KS.

## Acknowledgements

We are thankful for statistical analysis by Dr. Liliana Florea and Corina Antonescu, through the Computational Biology Consulting Core and support by Dr. Pletnikov in the JHMI behavioral core. H.T.B. is funded by the following sources: NIH: DP5OD017877, Louma G. Foundation, Icelandic Research Fund, #195835-051, #206806-051) and for this particular project with a grant from Takeda Pharmaceuticals.

## Data availability

All data have been posted to GEO and is available using accession numbers GSE146727, GSE146728, GSE146729.

