## Supplemental tables for "Inhibition of KDM1A activity restores adult neurogenesis and improves hippocampal memory in a mouse model of Kabuki syndrome"

### Table S1. Differentially bound H3K4me1 peaks from comparison of *Kmt2d*$^{+/\beta Geo}$ mice treated with vehicle versus *Kmt2d*$^{+/+}$ mice treated with vehicle

| Chr | Start | End | Distance to TSS | Fold change | *P*-value | Symbol |
|---|---|---|---|---|---|---|
| **Upregulated** | | | | | | |
| chr1 | 24684051 | 24684350 | 5282 | 3.15 | 4.56E-08 | *Lmbrd1* |
| chr17 | 13587351 | 13587750 | -33456 | 3.04 | 3.73E-13 | *2700054A10Rik* |
| chr11 | 31922601 | 31922900 | -42883 | 2.98 | 4.82E-13 | *4930524B15Rik* |
| chr1 | 85528851 | 85529150 | 69810 | 2.57 | 3.05E-06 | *Sp110* |
| **Downregulated** | | | | | | |
| chr3 | 75492601 | 75492950 | 64077 | 0.22 | 1.71E-07 | *Pdcd10* |
| chr7 | 83179601 | 83179950 | 31305 | 0.29 | 1.04E-08 | *A530021J07Rik* |
| chr4 | 1.46E+08 | 1.46E+08 | 16203 | 0.29 | 1.98E-05 | *Zfp980* |
| chr2 | 1.62E+08 | 1.62E+08 | -275288 | 0.30 | 3.48E-07 | *Ptprtos* |
| chr16 | 67016151 | 67016500 | 359207 | 0.32 | 1.64E-11 | *1700010K23Rik* |
| chr1 | 85180451 | 85180750 | -19011 | 0.32 | 1.72E-06 | *Gm7609* |
| chr2 | 1.34E+08 | 1.34E+08 | 128852 | 0.33 | 2.81E-06 | *Hao1* |
| chr3 | 69799701 | 69800050 | 60019 | 0.34 | 2.1E-08 | *Sptssb* |
| chr12 | 44648151 | 44648500 | 319440 | 0.34 | 6.68E-10 | *Nrcam* |
| chr2 | 1.77E+08 | 1.77E+08 | -25325 | 0.34 | 6.84E-06 | *Gm14295* |
| chr11 | 1.1E+08 | 1.1E+08 | -9049 | 0.35 | 1.48E-14 | *Abca6* |
| chr16 | 39393901 | 39394250 | 491730 | 0.36 | 8.15E-09 | *Igsf11* |
| chr18 | 19039551 | 19039850 | 962397 | 0.36 | 3.8E-06 | *Dsc3* |
| chr15 | 36567701 | 36568000 | -12278 | 0.38 | 9.4E-08 | *Snx31* |
| chr2 | 1.24E+08 | 1.24E+08 | -269444 | 0.39 | 4.97E-10 | *Sema6d* |
| chr4 | 1.48E+08 | 1.48E+08 | 3533 | 0.39 | 0.000696 | *Zfp979* |
| chr11 | 39221751 | 39222050 | 168325 | 0.39 | 2.27E-08 | *4930553C11Rik* |
| chr1 | 85238501 | 85238800 | 31916 | 0.39 | 0.000908 | *C130026I21Rik* |
| chr7 | 1.23E+08 | 1.23E+08 | 55509 | 0.39 | 3.19E-05 | *4930413G21Rik* |
| chr12 | 1.06E+08 | 1.06E+08 | -43909 | 0.40 | 1.8E-12 | *1700121N20Rik* |
| chr19 | 53964301 | 53964650 | 19599 | 0.40 | 8.38E-07 | *Shoc2* |
| chr17 | 16379951 | 16380250 | -553514 | 0.40 | 8.38E-10 | *Rgmb* |
| chr11 | 46955651 | 46956000 | 145026 | 0.40 | 2.37E-05 | *Timd4* |
| chr2 | 82344951 | 82345300 | 291467 | 0.41 | 6.32E-14 | *Zfp804a* |

### Table S2. Differentially bound H3K4me1 peaks upon comparing *Kmt2d*$^{+/\beta Geo}$ mutant mice treated with TAK-418 versus *Kmt2d*$^{+/+}$ mice treated with vehicle

| Chr | Start | End | Distance to TSS | Fold change | *P*-value | Description |
|---|---|---|---|---|---|---|
| **Upregulated** | | | | | | |
| chr15 | 38308451 | 38308800 | -7914 | 4.580961 | 1.09E-13 | *Klf10* |
| chr6 | 47757851 | 47758150 | -5075 | 4.23132 | 2.01E-08 | *Rn4.5s* |
| chr1 | 24684051 | 24684350 | 5282 | 3.949866 | 6.11E-09 | *Lmbrd1* |
| chr6 | 47672201 | 47672550 | 24 | 3.767908 | 2.43E-10 | *Rn4.5s* |
| chr6 | 1.16E+08 | 1.16E+08 | -27345 | 3.241102 | 3.45E-10 | *Plxnd1* |
| chr6 | 4923851 | 4924250 | 20730 | 3.084724 | 1.32E-08 | *Ppp1r9a* |
| chr8 | 39152901 | 39153300 | -144880 | 2.902813 | 1.98E-07 | *Gm6213* |
| chr1 | 1.71E+08 | 1.71E+08 | 602 | 2.835173 | 2.93E-08 | *Mir6546* |
| chrX | 1.7E+08 | 1.7E+08 | -17458 | 2.834468 | 8.93E-17 | *G530011O06Rik* |
| chr1 | 69456651 | 69457000 | 229135 | 2.756085 | 5.28E-08 | *Ikzf2* |
| chr1 | 88257601 | 88258050 | 11583 | 2.689857 | 3.49E-07 | *6430706D22Rik* |
| chr14 | 1.23E+08 | 1.23E+08 | 34747 | 2.478949 | 5.06E-05 | *Pcca* |
| chr17 | 13305901 | 13306450 | -48397 | 2.451861 | 1.63E-51 | *Tcp10c* |
| chrX | 1.7E+08 | 1.7E+08 | -26258 | 2.40602 | 5.48E-33 | *G530011O06Rik* |
| chr1 | 1E+08 | 1E+08 | 407535 | 2.316954 | 6.16E-05 | *Cntnap5b* |
| chrX | 1.7E+08 | 1.7E+08 | -15508 | 2.311163 | 9.46E-27 | *G530011O06Rik* |
| chr4 | 3176851 | 3177150 | 4917 | 2.299987 | 2.06E-06 | *Vmn1r2* |
| chr1 | 88286301 | 88287250 | -9196 | 2.294952 | 1.11E-10 | *Hjurp* |
| chr18 | 82170301 | 82170950 | -178545 | 2.293223 | 8.63E-15 | *Mir5127* |
| chr4 | 1.47E+08 | 1.47E+08 | 99559 | 2.250408 | 3.67E-06 | *Zfp982* |
| chr16 | 3239551 | 3239950 | -352648 | 2.226777 | 6.48E-06 | *Olfr161* |
| chr4 | 1.22E+08 | 1.22E+08 | -500850 | 2.217059 | 2.41E-05 | *Gm12887* |
| chr17 | 57855901 | 57856300 | 86530 | 2.144734 | 8.81E-11 | *Cntnap5c* |
| **Downregulated** | | | | | | |
| chr1 | 85463551 | 85463850 | 135110 | 0.462643 | 4.47E-05 | *Sp110* |
| chr4 | 1.47E+08 | 1.47E+08 | -128525 | 0.444317 | 0.000501 | *Rex2* |
| chr1 | 85010801 | 85011150 | -75892 | 0.436585 | 6.64E-05 | *Slc16a14* |
| chrX | 1.27E+08 | 1.27E+08 | 140604 | 0.425518 | 2.04E-09 | *4932411N23Rik* |
| chr1 | 85312151 | 85312500 | -41759 | 0.419504 | 0.000288 | *C130026I21Rik* |
| chr8 | 95117501 | 95117800 | -4308 | 0.413902 | 2.52E-05 | *Kifc3* |
| chr8 | 20382251 | 20382600 | -19223 | 0.398451 | 1.19E-05 | *Gm15319* |

### Table S3. Differentially bound H3K4me3 peaks upon comparison of *Kmt2d*$^{+/\beta Geo}$ on vehicle versus *Kmt2d*$^{+/+}$ mice treated with vehicle.

| Symbol | Chr | Start | End | *P*-value | Fold change | Description |
|---|---|---|---|---|---|---|
| **Upregulated** | | | | | | |
| *Cnot9* | chr1 | 74506301 | 74506800 | 5.83E-06 | 2.750 | CCR4-NOT transcription complex, subunit 9 |
| *Zfp800* | chr6 | 28261001 | 28261350 | 5.95E-04 | 2.500 | zinc finger protein 800 |
| **Downregulated** | | | | | | |
| *Tspan2* | chr3 | 1.03E+08 | 1.03E+08 | 1.73E-07 | 0.258 | tetraspanin 2 |
| *B3gnt5* | chr16 | 19761051 | 19761400 | 1.83E-06 | 0.264 | UDP-GlcNAc:betaGal beta-1,3-N-acetylglucosaminyltransferase 5 |
| *Zbtb18* | chr1 | 1.77E+08 | 1.77E+08 | 4.31E-07 | 0.271 | zinc finger and BTB domain containing 18 |
| *4833420G17Rik* | chr13 | 1.19E+08 | 1.19E+08 | 2.53E-02 | 0.280 | RIKEN cDNA 4833420G17 gene |
| *Ddx3x* | chrX | 13281151 | 13281800 | 2.54E-12 | 0.299 | DEAD/H (Asp-Glu-Ala-Asp/His) box polypeptide 3, X-linked |
| *Zfp503* | chr14 | 21987651 | 21987750 | 1.23E-02 | 0.315 | zinc finger protein 503 |
| *Asns* | chr6 | 7692901 | 7693200 | 1.05E-06 | 0.318 | asparagine synthetase |
| *Mfap1b* | chr2 | 1.21E+08 | 1.21E+08 | 2.46E-05 | 0.319 | microfibrillar-associated protein 1B |
| *Cfap36* | chr11 | 29247101 | 29247400 | 4.37E-05 | 0.321 | cilia and flagella associated protein 36 |
| *Nfkbiz* | chr16 | 55821701 | 55822050 | 9.65E-06 | 0.323 | nuclear factor of kappa light polypeptide gene enhancer in B cells inhibitor, zeta |
| *Osbpl6* | chr2 | 76406201 | 76406400 | 2.10E-03 | 0.323 | oxysterol binding protein-like 6 |
| *Tubb4a* | chr17 | 57087351 | 57087500 | 3.21E-03 | 0.327 | tubulin, beta 4A class IVA |
| *Rerg* | chr6 | 1.37E+08 | 1.37E+08 | 4.38E-04 | 0.327 | RAS-like, estrogen-regulated, growth-inhibitor |
| *Pagr1a* | chr7 | 1.27E+08 | 1.27E+08 | 1.37E-04 | 0.332 | PAXIP1 associated glutamate rich protein 1A |
| *Cacfd1* | chr2 | 27010051 | 27010400 | 3.73E-06 | 0.341 | calcium channel flower domain containing 1 |
| *Tmem151b* | chr17 | 45549251 | 45549650 | 1.60E-05 | 0.342 | transmembrane protein 151B |
| *Hebp1* | chr6 | 1.35E+08 | 1.35E+08 | 2.36E-04 | 0.347 | heme binding protein 1 |
| *Tmem55a* | chr4 | 14864301 | 14864750 | 1.73E-07 | 0.348 | transmembrane protein 55A |
| *Narf* | chr11 | 1.21E+08 | 1.21E+08 | 6.89E-06 | 0.348 | nuclear prelamin A recognition factor |
| *BC055324* | chr1 | 1.64E+08 | 1.64E+08 | 7.22E-04 | 0.348 | cDNA sequence BC055324 |
| *Akap1* | chr11 | 88864201 | 88864600 | 1.06E-06 | 0.349 | A kinase (PRKA) anchor protein 1 |
| *Etv1* | chr12 | 38778801 | 38779250 | 2.76E-05 | 0.350 | ets variant 1 |

**Table S4. Differentially bound H3K4me3 peaks upon comparison of *Kmt2d*$^{+/\beta Geo}$ mice on TAK-418 versus *Kmt2d*$^{+/+}$ mice treated with vehicle.**

| Symbol | Chr | Start | End | P-value | Fold change | Gene.Description |
|---|---|---|---|---|---|---|
| **Upregulated** | | | | | | |
| *Col19a1* | chr1 | 24613301 | 24613950 | 4.99E-17 | 4.091 | collagen, type XIX, alpha 1 |
| *Ajap1* | chr4 | 1.53E+08 | 1.53E+08 | 2.81E-14 | 4.050 | adherens junction associated protein 1 |
| *6820431F20Rik* | chr8 | 20150551 | 20150950 | 4.31E-08 | 3.538 | cadherin 11 pseudogene |
| **Downregulated** | | | | | | |
| *Ddx6* | chr9 | 44605601 | 44606000 | 3.11E-07 | 0.172 | DEAD (Asp-Glu-Ala-Asp) box polypeptide 6 |
| *B430212C06Rik* | chr18 | 67343651 | 67344100 | 9.01E-10 | 0.188 | RIKEN cDNA B430212C06 gene |
| *2310022B05Rik* | chr8 | 1.25E+08 | 1.25E+08 | 9.21E-11 | 0.200 | RIKEN cDNA 2310022B05 gene |
| *1810014B01Rik* | chr10 | 86685501 | 86685950 | 9.46E-09 | 0.202 | RIKEN cDNA 1810014B01 gene |
| *Smad7* | chr18 | 75342701 | 75343050 | 4.97E-07 | 0.207 | SMAD family member 7 |
| *Mri1* | chr8 | 84256851 | 84257250 | 1.57E-07 | 0.208 | methylthioribose-1-phosphate isomerase 1 |
| *Rela* | chr19 | 5637651 | 5638050 | 1.89E-07 | 0.211 | v-rel reticuloendotheliosis viral oncogene homolog A (avian) |
| *Trim59* | chr3 | 69044301 | 69044800 | 5.31E-11 | 0.212 | tripartite motif-containing 59 |
| *Lca5l* | chr16 | 96191901 | 96192300 | 7.84E-07 | 0.221 | Leber congenital amaurosis 5-like |
| *Vapb* | chr2 | 1.74E+08 | 1.74E+08 | 6.89E-09 | 0.229 | vesicle-associated membrane protein, associated protein B and C |
| *Pskh1* | chr8 | 1.06E+08 | 1.06E+08 | 2.49E-06 | 0.232 | protein serine kinase H1 |
| *Parp4* | chr14 | 56575501 | 56575850 | 6.95E-05 | 0.238 | poly (ADP-ribose) polymerase family, member 4 |
| *Suclg2* | chr6 | 95718501 | 95718850 | 1.37E-06 | 0.239 | succinate-Coenzyme A ligase, GDP-forming, beta subunit |
| *Mfap3l* | chr8 | 60632351 | 60632700 | 1.92E-05 | 0.239 | microfibrillar-associated protein 3-like |
| *Patl1* | chr19 | 11912601 | 11913150 | 3.19E-07 | 0.241 | protein associated with topoisomerase II homolog 1 (yeast) |
| *Tmem120b* | chr5 | 1.23E+08 | 1.23E+08 | 1.96E-05 | 0.242 | transmembrane protein 120B |
| *Them6* | chr15 | 74721351 | 74721800 | 5.89E-07 | 0.247 | thioesterase superfamily member 6 |
| *Ptov1* | chr7 | 44869301 | 44869700 | 4.36E-06 | 0.253 | prostate tumor over expressed gene 1 |
| *Zfp334* | chr2 | 1.65E+08 | 1.65E+08 | 5.89E-06 | 0.255 | zinc finger protein 334 |
| *Mir6921* | chr11 | 60176651 | 60177250 | 5.64E-07 | 0.255 | microRNA 6921 |
| *Dcaf5* | chr12 | 80434951 | 80435350 | 8.47E-08 | 0.256 | DDB1 and CUL4 associated factor 5 |
| *Slc1a2* | chr2 | 1.03E+08 | 1.03E+08 | 6.06E-06 | 0.257 | solute carrier family 1 (glial high affinity glutamate transporter), member 2 |
| *Zbtb43* | chr2 | 33468101 | 33468500 | 2.43E-06 | 0.258 | zinc finger and BTB domain containing 43 |
| *Exosc9* | chr3 | 36552651 | 36553000 | 6.43E-05 | 0.259 | exosome component 9 |

### Table S5. Differentially expressed genes upon comparison of *Kmt2d*$^{+/\beta Geo}$ mice on vehicle versus *Kmt2d*$^{+/+}$ mice on vehicle.

| Symbol | Description | Fold change | *P* value |
|---|---|---|---|
| **Upregulated** | | | |
| *Nlrp5-ps* | NLR family, pyrin domain containing 5, pseudogene | 2.319 | 7.69E-05 |
| *Onecut1* | one cut domain, family member 1 | 2.095 | 4.82E-04 |
| *Fmod* | Fibromodulin | 1.734 | 1.75E-04 |
| *A4galt* | alpha 1,4-galactosyltransferase | 1.722 | 2.41E-02 |
| *Clec3b* | C-type lectin domain family 3, member b | 1.719 | 7.90E-03 |
| *Ptx3* | pentraxin related gene | 1.642 | 1.83E-02 |
| *Mlph* | Melanophilin | 1.638 | 2.10E-02 |
| *Xdh* | xanthine dehydrogenase | 1.636 | 1.23E-02 |
| *Kcnh6* | potassium voltage-gated channel, subfamily H (eag-related), member 6 | 1.601 | 4.75E-10 |
| *Eln* | Elastin | 1.587 | 1.71E-07 |
| *F13a1* | coagulation factor XIII, A1 subunit | 1.575 | 2.59E-02 |
| **Downregulated** | | | |
| *Trh* | thyrotropin releasing hormone | 0.112 | 1.85E-02 |
| *Rxfp1* | relaxin/insulin-like family peptide receptor 1 | 0.448 | 5.10E-08 |
| *Nxph3* | neurexophilin 3 | 0.469 | 3.92E-06 |
| *Tshz2* | teashirt zinc finger family member 2 | 0.484 | 1.94E-04 |
| *Plcz1* | phospholipase C, zeta 1 | 0.500 | 8.14E-04 |
| *Fos* | FBJ osteosarcoma oncogene | 0.535 | 4.39E-02 |
| *Egr2* | early growth response 2 | 0.563 | 8.96E-03 |
| *Olfr287* | olfactory receptor 28 | 0.567 | 1.79E-05 |
| *Susd2* | sushi domain containing 2 | 0.577 | 3.78E-03 |
| *Nts* | Neurotensin | 0.577 | 1.48E-02 |
| *Nhlh1* | nescient helix loop helix 1 | 0.607 | 1.24E-02 |
| *Cbln1* | cerebellin 1 precursor protein | 0.625 | 8.97E-03 |
| *Otof* | Otoferlin | 0.627 | 2.46E-04 |
| *Olfr55* | olfactory receptor 55 | 0.668 | 4.30E-02 |
| *Ntn5* | netrin 5 | 0.683 | 1.34E-02 |
| *Bub1b* | mitotic checkpoint serine/threonine kinase | 0.683 | 3.62E-03 |
| *Nov* | nephroblastoma overexpressed gene | 0.685 | 1.12E-02 |
| *Igfbp6* | insulin-like growth factor binding protein 6 | 0.688 | 2.46E-03 |
| *Met* | met proto-oncogene | 0.690 | 2.67E-03 |
| *Fosb* | FBJ osteosarcoma oncogene B | 0.696 | 4.73E-03 |
| *Arc* | activity regulated cytoskeletal-associated protein | 0.730 | 3.76E-02 |
| *Junb* | jun B proto-oncogene | 0.737 | 2.82E-04 |
| *Kcnf1* | potassium voltage-gated channel, subfamily F, member 1 | 0.755 | 9.99E-09 |

### Table S6. Differentially expressed genes upon comparison of *Kmt2d*$^{+/\beta Geo}$ mice on TAK-418 versus *Kmt2d*$^{+/+}$ mice on vehicle control.

| Symbol | Description | Fold change | *P*-value |
|---|---|---|---|
| **Upregulated** | | | |
| *Prph* | Peripherin | 3.040 | 7.04E-05 |
| *Atp4a* | ATPase, H+/K+ exchanging, gastric, alpha polypeptide | 2.226 | 6.27E-04 |
| *Lefty2* | left-right determination factor 2 | 2.131 | 3.16E-05 |
| *Nlrp5-ps* | NLR family, pyrin domain containing 5, pseudogene | 1.953 | 1.51E-03 |
| *Gipc3* | GIPC PDZ domain containing family, member 3 | 1.865 | 1.02E-04 |
| *Espn* | Espin | 1.863 | 4.17E-10 |
| *Arsi* | arylsulfatase i | 1.788 | 4.62E-03 |
| *Onecut1* | one cut domain, family member 1 | 1.740 | 5.63E-03 |
| *Samd11* | sterile alpha motif domain containing 11 | 1.710 | 2.11E-02 |
| *Ppp1r32* | protein phosphatase 1, regulatory subunit 32 | 1.692 | 6.83E-03 |
| *Slc16a3* | solute carrier family 16 (monocarboxylic acid transporters), member 3 | 1.674 | 1.55E-03 |
| *Tnxb* | tenascin XB | 1.658 | 3.26E-14 |
| *Duxbl3* | double homeobox B-like 3 | 1.636 | 1.84E-02 |
| *Arhgap36* | Rho GTPase activating protein 36 | 1.630 | 8.82E-04 |
| *Rps6kc1* | ribosomal protein S6 kinase polypeptide 1 | 1.617 | 4.43E-04 |
| *Plp2* | proteolipid protein 2 | 1.615 | 2.60E-03 |
| *Lrrn4* | leucine rich repeat neuronal 4 | 1.613 | 1.72E-02 |
| **Downregulated** | | | |
| *Trh* | thyrotropin releasing hormone | 0.167 | 4.40E-02 |
| *Baiap2l1* | BAI1-associated protein 2-like 1 | 0.362 | 6.94E-04 |
| *Egr2* | early growth response 2 | 0.550 | 1.02E-03 |
| *Cd59a* | CD59a antigen | 0.612 | 1.65E-03 |
| *Otof* | Otoferlin | 0.627 | 3.62E-04 |
| *Ppfibp2* | PTPRF interacting protein, binding protein 2 (liprin beta 2) | 0.628 | 4.86E-02 |
| *Mir129-1* | microRNA 129-1 | 0.636 | 1.66E-03 |
| *Rxfp1* | relaxin/insulin-like family peptide receptor 1 | 0.660 | 1.86E-03 |
| *Satb2* | special AT-rich sequence binding protein 2 | 0.669 | 1.74E-05 |
| *Iqub* | IQ motif and ubiquitin domain containing | 0.676 | 4.76E-02 |
| *Fam46c* | family with sequence similarity 46, member C | 0.678 | 2.25E-02 |
