## Supplemental figures for "Inhibition of KDM1A activity restores adult neurogenesis and improves hippocampal memory in a mouse model of Kabuki syndrome"

**Supplementary Figures**

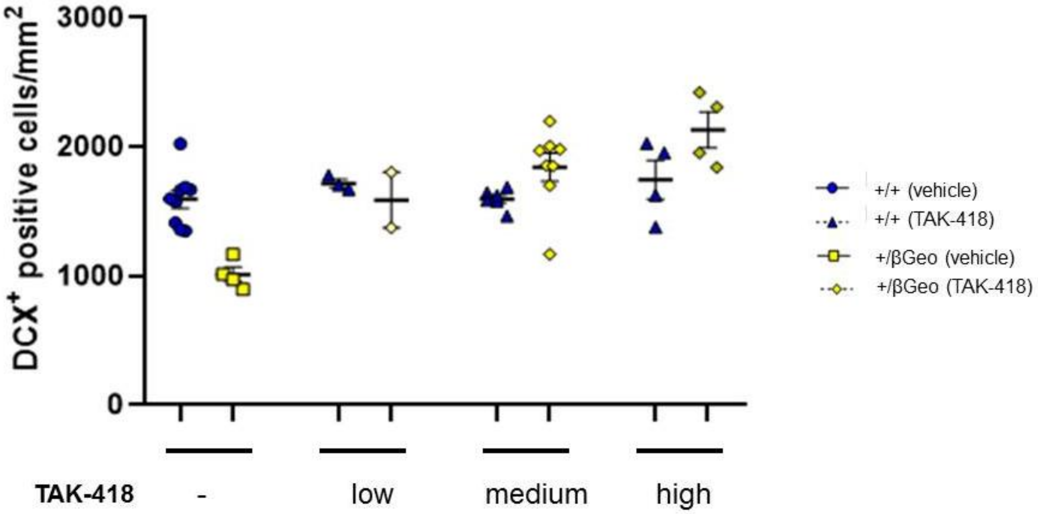


**Supplementary Fig. 1. TAK-418 demonstrates dose-dependent rescue of the number of DCX$^+$ cells.** Both genotypes were treated with TAK-418 and a dose dependent response on this metric in *Kmt2d*$^{+/\beta Geo}$ mice was observed with no obvious effect on *Kmt2d*$^{+/+}$ littermates.

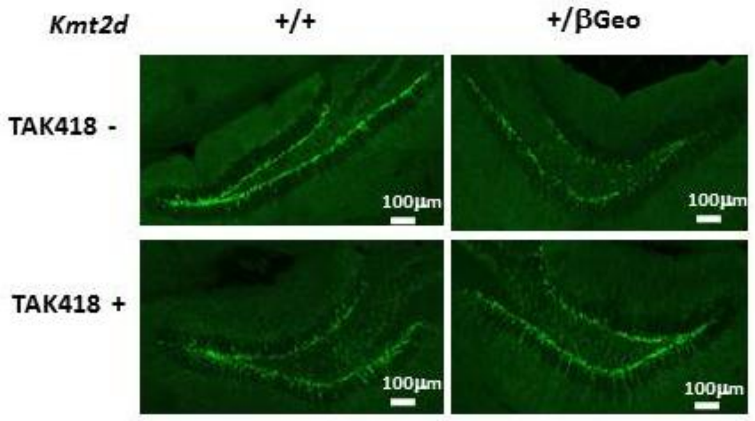


**Supplementary Fig. 2. Representative images of the dentate gyrus after immunofluorescence staining for Doublecortin.** On visual inspection the $DCX^{+}$ processes appeared shorter in the *Kmt2d*$^{+/\beta Geo}$ mice compared to wildtype littermates but longer in the *Kmt2d*$^{+/\beta Geo}$ mice on TAK-418.

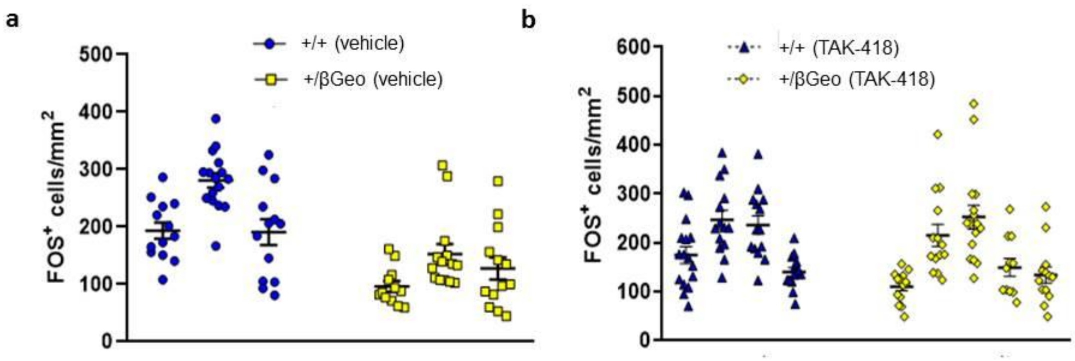


**Supplementary Fig. 3. Quantification of FOS$^+$ cells in the dentate gyrus area of hippocampus after immunofluorescent staining against FOS.** Each point represents total FOS$^+$ cells from one brain slice. The mouse genotype and treatment is indicated. These data were averaged for figure 4 (**f, g**).

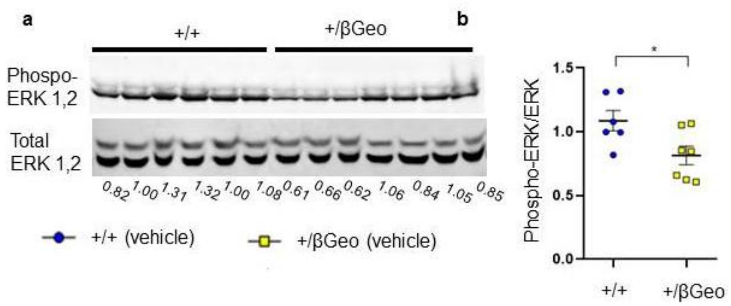


**Supplementary Fig. 4. Hippocampal extracts from *Kmt2d*$^{+/\beta Geo}$ mice reveal a deficiency of ERK activation compared to littermates.** Western blots from *Kmt2d*$^{+/\beta Geo}$ mice and wildtype littermates reveal a decrease in phosphoERK/ERK **(a, b).**

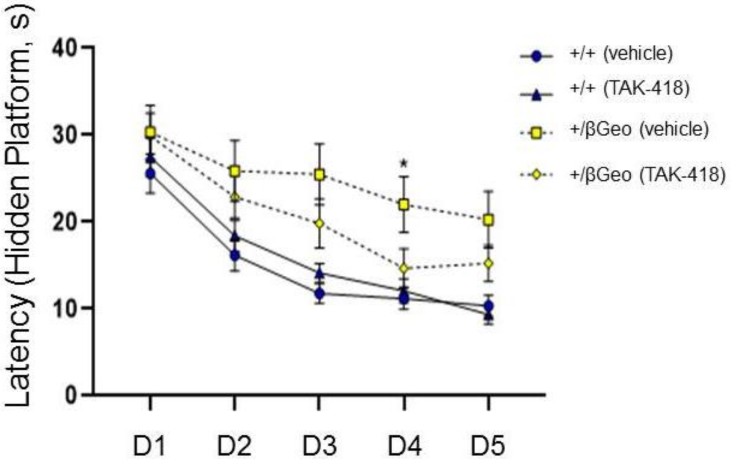


**Supplementary figure 5. The *Kmt2d*$^{+/\beta Geo}$ mice demonstrate a trend toward increased latency to reach the hidden platform compared to wildtype littermates which was only significant on day 4.** These data represent our hidden platform training over 5 individual days (D). Each group represents 12-14 mice. *$p < 0.05$.

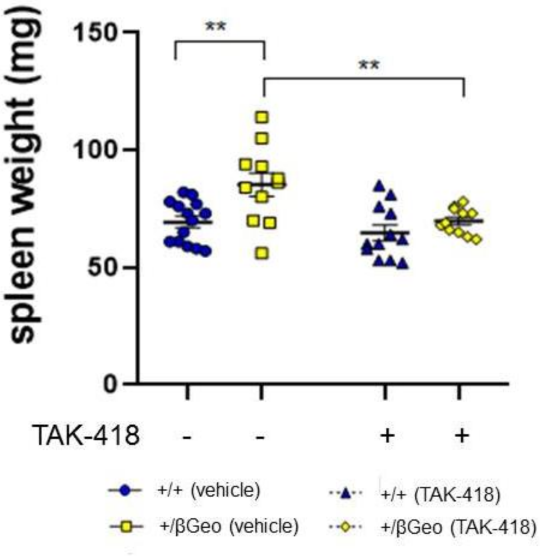


**Supplementary Fig. 6. Splenomegaly normalizes in the *Kmt2d*$^{+/\beta Geo}$ mice treated with TAK-418 for 2 months.** Treatment was initiated at 2 months of age. Spleens were weighed at the time of sacrifice. Each point represents one mouse; the genotype of the mice and treatment received is indicated. **p < 0.01.
